# Accurate single-bead force calibration in high-throughput magnetic tweezers reveals the mechanism of directional transcription termination by MTERF1

**DOI:** 10.64898/2026.04.17.719150

**Authors:** Pim P. B. America, Eugeniu Ostrofet, Britney Johnson, Salina Quack, Flavia Stal-Papini, Quinte Smitskamp, Daniel Buc, Jamie J. Arnold, Craig E. Cameron, David Dulin

**Affiliations:** Department of Physics and Astronomy, and LaserLaB Amsterdam, Vrije Universiteit Amsterdam, De Boelelaan 1081, 1081 HV, Amsterdam, the Netherlands; Junior Research Group 2, Interdisciplinary Center for Clinical Research, Friedrich-Alexander-University Erlangen-Nürnberg (FAU), Cauerstr. 3, 91058 Erlangen, Germany; Department of Microbiology and Immunology, University of North Carolina School of Medicine, Chapel Hill, NC 27599, USA

## Abstract

High-throughput force spectroscopy assays, such as with magnetic tweezers, enable reconstruction of biomolecular reaction energy landscapes and provide access to rare events with deep statistics. Precise force calibration is essential for accurately describing complex reactions, which can be hindered by sample heterogeneity, such as bead-to-bead difference in magnetic content. Here, we describe an *in-situ* force calibration methodology for high-throughput magnetic tweezers that enables the calibration for each individual bead with an accuracy of up to 3%, limited only by the statistical resolution. We apply this approach to characterize the directional transcription termination molecular mechanism by the polar roadblock mitochondrial transcription termination factor 1 (MTERF1). Establishing a SpyTag-SpyCatcher surface-attachment strategy, we performed force-jump experiments on the same tethers for up to 11 hours. We showed that directional DNA unwinding is sufficient to explain the polar roadblock activity of MTERF1. Accurate force spectroscopy further reveals that the unlocking transition is rate-limited by a single kinetic barrier, with a transition-state distance consistent with structural interpretations. Together, these results provide a mechanistic and broadly applicable model for the asymmetric stability of MTERF1 and other nucleic acid polar roadblocks and establish a robust force spectroscopy framework for high-throughput magnetic tweezers experiments.

## INTRODUCTION

Single-molecule force spectroscopy techniques enable the physical manipulation of biomolecular reactions, allowing precise interrogation of their energy landscape (1). They are separated in two main categories, position clamp and force clamp. The former includes optical tweezers and atomic force microscopy (AFM), and the latter involved techniques such as acoustic force spectroscopy (AFS) (2), centrifuge force spectroscopy (3), flow stretching and magnetic tweezers (4).

The latter are well suited for multiplexed, high-throughput force spectroscopy experiments (5), and have become increasingly popular for their ability to rapidly collect a large number of time-traces on complex biomolecular reactions, capturing even the rarest event in great detail (6). Near equilibrium force spectroscopy measurements for day-long experiments have been demonstrated with magnetic tweezers (7). High-throughput magnetic tweezers in combination with force-jump experiments have been employed to characterize the molecular mechanism behind directional *E. coli* bacterial replication termination by the Tus-*Ter* complex (8). Recent developments in DNA and protein scaffolds in combination with magnetic tweezers and AFS have enabled force-jump experiments on the same tether over many cycles and for many tethers in parallel, enabling the precise interrogation of ligand-ligand interactions under force (9–13). These assays require precise force calibration to enable accurate reconstruction of the energy landscape for these biomolecular reactions.

Force heterogeneity in high-throughput single-molecule force spectroscopy can arise from two different factors: inhomogeneity in the force field, e.g. with AFS (14), or in the force transducer, e.g. magnetic material content in magnetic beads used in magnetic tweezers (15– 17). Such inhomogeneity makes force calibration standard, i.e. force calibration obtained from averaging force measurements over many beads in the field of view (16, 17), not accurate enough to enable precise energy landscape reconstruction. Several methods have been reported for magnetic tweezers force calibration (15–19), which are mainly adapted for measurements on long nucleic acid tethers (several microns long) and are not adapted for force calibration using short tethers. A power spectrum approach has been used to address this problem (20), but such approach may be too difficult to implement for non-physicists. A simple and robust – yet accurate – force calibration method would therefore be of interest for a broader, non-specialist user-base.

The human mitochondrial genome is a gene-dense, ∼16.6 kbp long polycistronic plasmid that encodes for many genes on both strands, called heavy and light strand (HS and LS) (21) (**Fig. 1A**). The transcription of both strands is performed by the mitochondrial RNA polymerase (mtRNAP) that initiates RNA synthesis in the non-coding region (NCR, **Fig. 1A**) from either the heavy or the light strand promoter (HSP and LSP, respectively), leading to bidirectional transcription (21) (**Fig. 1A**). LS transcription termination is mediated by the mitochondrial transcription termination factor 1 (MTERF1) bound to a 21 bp termination site immediately downstream of the 16S rRNA gene encoded on the HS (22, 23), while HS transcribing mtRNAP are only partially hindered (**Fig. 1A**) (24, 25). Structural and functional studies have highlighted how MTERF1 binding with the termination site results in local DNA unwinding and finally a locked complex (26–28). MTERF1 residues form specific stacking interactions with three everted bases, i.e. A3243 (Tyr288), T3243 (Arg162) and C3242 (Phe243) (**Fig. 1B**) (27, 29). Mutating any of these residues, such as Arg162, results in a loss of transcription termination efficiency (27, 29). MTERF1 is also found to pause the elongating mitochondrial replisome *in vivo* by acting as a barrier to the mitochondrial helicase TWINKLE (30, 31). MTERF1 has therefore been proposed to regulate the dynamics of mitochondrial transcription and replication to avoid RNAP–RNAP or RNAP–replisome heads-on collisions. Despite numerous structural and biochemical studies, the molecular mechanism behind the MTERF1 polar roadblock activity remains unclear.

**Figure 1:**
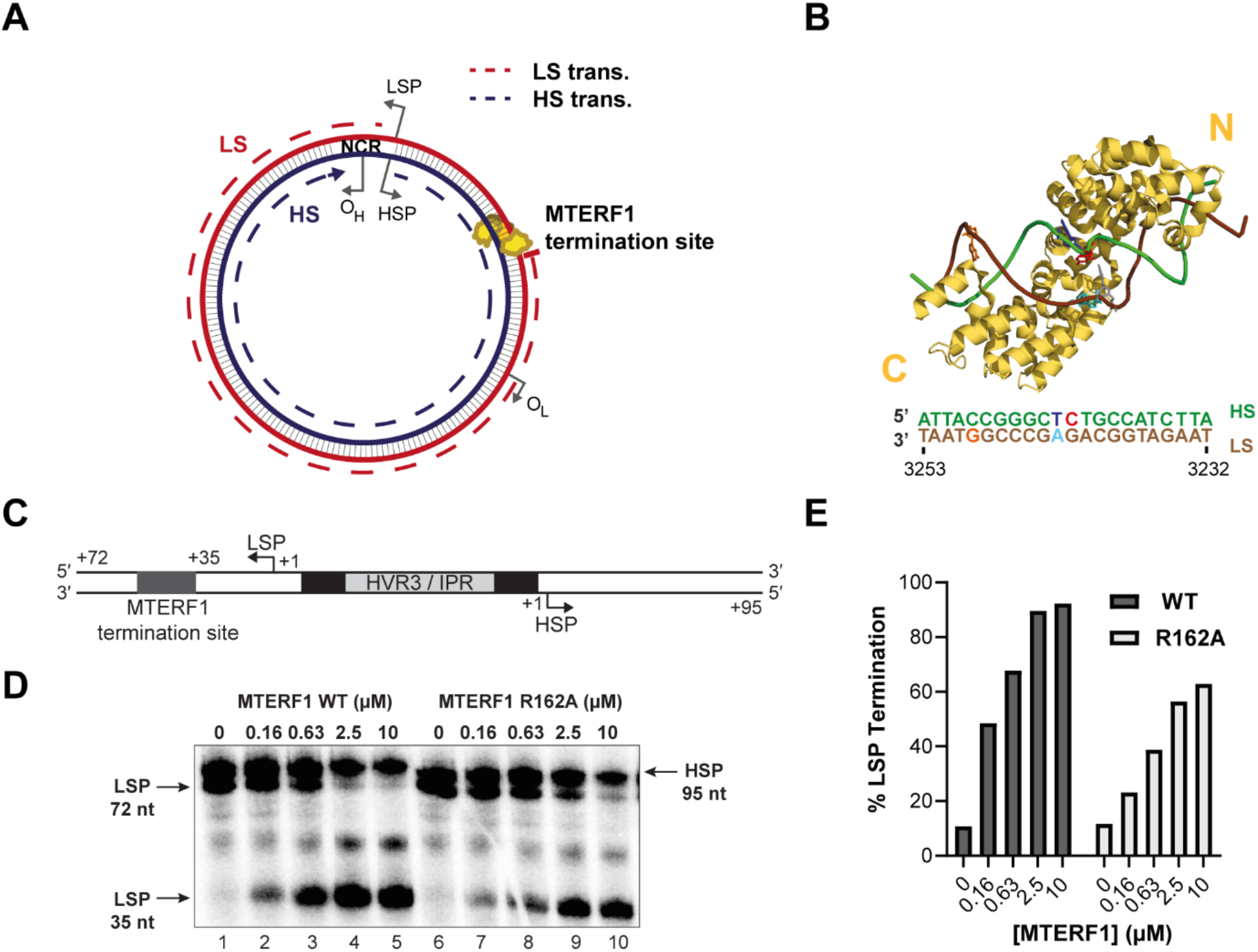
MTERF1 terminates LS-transcribing human mitochondrial RNA polymerase. **(A)** Illustration of the human mitochondrial genome, with the light strand (LS) on the outer side (red solid line) and the heavy strand (HS) on the inner side of the mitochondrial genome representation (blue solid line). LSP and HSP represent the light and heavy strand promoters, O_H_ and O_L_ denote the origins of replication. Blue and red dashed circular arrows indicate the direction of transcription for HS and LS strands, respectively. MTERF1 is represented bound to LS transcription termination site. **(B)** Ribbon representation of MTERF1 structure (the N- and C-terminus indicated by yellow N and C) bound to the termination site (sequence is indicated below with the same color code). In the bound complex C3242, A3243 and T3243 are everted and stacked, stabilizing the complex. In the A3243G mutant, the A-T base pair (light and dark blue) is replaced by a G-C. The G3249A base pair mutation is indicated in orange. Adapted from PDB accessing number: 3MVA. **(C)** DNA construct design used for the *in vitro* transcription experiments. **(D)** Run-off transcription products from in vitro mitochondrial transcription experiment using the construct described in (C) for different concentrations of either wild-type (WT) or Arginine 162 (R162A) mutant MTERF1, resolved by denaturing PAGE. **(E)** Percentage (%) of terminated transcription product from the LS promoter (LSP 35 nt relative to 72 nt) for different concentrations of MTERF1 for either WT (grey bars) or the R162A mutation (white bars) quantified from (D).

Here, we investigated the mechanism of MTERF1 directional transcription termination. Bulk assays show that MTERF1 enables efficient transcription termination of LS-transcribing mtRNAPs, and MTERF1 R162A mutation significantly impairs termination efficiency. We developed a single-molecule high-throughput magnetic tweezers assay to performed force-jump experiments on DNA hairpins encoding a termination site in either orientation, enabling directional DNA unwinding towards either the C- or N-terminus of MTERF1 (**Fig. 1B**). We found that MTERF1 has an asymmetric stability upon directional DNA unwinding, with the non-permissive orientation providing the most stable lock. We further show that the stability as a function of the stacking interactions between bases on the termination site and specific MTERF1 binding residues, such as R162, suggesting that the lock stability under directional DNA unwinding is important for directional mtRNAP transcription termination. We established an *in-situ* force calibration for high-throughput magnetic tweezers, enabling accurate force spectroscopy – even for short tethers – that is only limited by the statistical resolution. We also implemented a tether attachment strategy using a SpyTag-SpyCatcher approach (32, 33) to increase the tether surface attachment stability with regard to the commonly used digoxigenin– anti-digoxigenin attachment. This strategy enabled force-jump experiments on the same tethers for more than eleven hours. Using these developments, we found that MTERF1 unlocking in the non-permissive orientation is rate-limited by a single kinetic transition, and the distance to the transition state is consistent with the spacing of the three everted bases. We propose a mechanistic model describing MTERF1 lock origin on the termination that is potentially applicable to other polar roadblocks on nucleic acids. We anticipate that our approach combining high-throughput magnetic tweezers and accurate force spectroscopy measurements will be broadly applicable to precisely measure distances in the energy landscape of complex biomolecular reactions.

## MATERIALS AND METHODS

### DNA hairpin fabrication with biotin and digoxygenin handles

The fabrication of the DNA hairpins with biotin and digoxigenin has been described in detail by (34). The DNA hairpin used in this study (**Table S1A**) is made of a 1049 bp double-stranded DNA stem terminated by a ∼10 nt loop and two handles of 432 bp at the 3′-end and 951 bp at the 5′-end that is assembled from nine ssDNA oligos, which are PCR amplified and annealed together. The stem is PCR amplified from a plasmid that contains a binding site for MTERF1 in either the permissive or non-permissive direction (**Table S1A**). The handles are PCR amplified from a λ1 DNA plasmid (35). These include a biotin-labelled ssDNA at the 3′-end and a digoxygenin-labelled ssDNA at the 5′-end to attach to the streptavidin-coated magnetic bead (M270) and to the anti-digoxigenin coated glass surface, respectively.

### DNA hairpin fabrication with biotin and SpyTag handles

The fabrication of the DNA hairpins has been described in detail by (34). The DNA hairpin used for the experiments of very long duration (**Table S1B**) is also made of a 1049 bp double-stranded DNA stem terminated by a ∼10 nt loop and two handles of 432 bp at the 3′-end and 519 bp at the 5′-end that is assembled from eight ssDNA oligos, which are PCR amplified and annealed together. The stem contains a binding site for MTERF1 in either the permissive or non-permissive direction. The handles include a biotin-labelled ssDNA at the 3′-end to attach to the streptavidin-coated magnetic bead (M270, DynaBeads) and an oligo-peptide conjugate of ssDNA with the SpyTag (Eurogentec) at the 5′-end to the Spycatcher3 (Bio-Rad) coated glass surface, respectively.

### RNA hairpin fabrication with biotin and digoxygenin handles

The fabrication of the RNA hairpins has also been described by (34). The RNA hairpins used for the proof of concept of the *in-situ* force calibration method is given in **Table S1C**. This hairpin has a 499 bp RNA stem and 4 nt loop. The two RNA handles are 842 bp with a biotin handle and 863 bp with a digoxigenin handle. The handles are obtained by PCR amplification on a λ1 DNA plasmid (35) with several primers and the hairpin stem is obtained from PCR amplification on a pMK-RQ plasmid (**Table S1C**). The PCR products are purified, in-vitro transcribed, annealed and eventually ligated (34). The digoxygenin-labeled ssRNA and the biotin-labeled ssRNA handles on either side are attached to the anti-digoxigenin coated glass surface and the streptavidin-coated magnetic bead (M-270), respectively.

### Long dsDNA tether fabrication with biotin and digoxigenin handles

The 20.6 kbp DNA construct used for the reference force calibration, is constructed from a λ1 DNA plasmid. The biotin and digoxigenin handles are obtained from PCR with the forward primer AAAAGCGGCCGCCCAGCGAGTCACTCAGCGC and reverse primer AAAACTCGAGTCTGCTGCTCAGCCTTC (ThermoFisher Scientific). Then the construct is obtained by digesting of the plasmid with the restriction enzymes *Not*I and *Xho*I, which are trim the DNA handles and create the long spacer. After the formation of the long dsDNA it is ligated with T4 DNA ligase (New England Biolabs) (17).

### High-throughput magnetic tweezers apparatus

The high-throughput magnetic tweezers used in this study have already been described in detail in (36). Briefly, two vertically aligned permanent magnets (5 mm cubes, SuperMagnete, Switzerland) separated by a 1 mm gap are positioned above a flow cell (see paragraph below) which is mounted on a custom-built inverted microscope. The vertical position and rotation of the magnets are controlled by two linear motors, M-126-PD1 and C-150 (Physik Instrumente PI, GmbH & Co. KG, Karlsruhe, Germany), respectively. The field of view is illuminated through the magnets gap by a collimated LED-light source and is imaged onto a large chip CMOS camera (Dalsa Falcon2 FA-80-12M1H, Stemmer Imaging, Germany) using a 50× oil immersion objective (CFI Plan Achro 50 XH, NA 0.9, Nikon, Germany) and an achromatic doublet tube lens of 200 mm focal length and 50 mm diameter (Qioptic, Germany). To control the temperature, we used a system described in details in (37). A flexible resistive foil heater with an integrated 10 MΩ thermistor (HT10K, Thorlabs) is wrapped around the microscope objective and further insulated by several layers Kapton tape (KAP22-075, Thorlabs). The heating foil is connected to a PID temperature controller (TC200 PID controller, Thorlabs) to adjust the temperature within ∼0.1 °C.

### Flow cell assembly with anti-digoxigenin surface

The fabrication procedure for flow cells has been described in detail in Ref. (36). To summarize, we sandwiched a double layer of Parafilm by two #1 coverslips, the top one having one hole at each end serving as inlet and outlet, the bottom one being coated with a 0.1% m/V nitrocellulose dissolved in amyl acetate solution. The flow cell was mounted into a custom-built holder and rinsed with ∼1 ml of 1x phosphate buffered saline (PBS). 3 μm diameter polystyrene reference beads were attached to the bottom coverslip surface by incubating 100 μl of a 1:1000 dilution in PBS (LB30, Sigma Aldrich) for ∼3 minutes. 50 μl of anti-digoxigenin (50 μg/ml in PBS) was incubated for 30 minutes in the flow cell. The flow cell was then flushed with 1 ml of 10 mM Tris, 1 mM EDTA pH 8.0, 750 mM NaCl, 2 mM sodium azide to remove excess of anti-digoxigenin followed by rinsing with another 0.5 ml of TE buffer (10 mM Tris, 1 mM EDTA pH 8.0, 150 mM NaCl, 2 mM sodium azide). The surface was then passivated with a 10 mg/ml solution of bovine serum albumin (BSA, Fraction V, Carl Roth) in PBS and 5% glycerol for 30 minutes, and rinsed with 0.5 ml of TE buffer.

### Flow cell assembly with cross-linked anti-digoxigenin surface

The fabrication procedure for flow cells has been described in detail in (36). To summarize, we sandwiched a double layer of Parafilm by two #1 coverslips, the top one having one hole at each end serving as inlet and outlet, the bottom one being coated with a 0.1% m/V nitrocellulose dissolved in amyl acetate solution. The flow cell was mounted into a custom-built holder and rinsed with ∼1 ml of 1x phosphate buffered saline (PBS). 3 μm diameter polystyrene reference beads were attached to the bottom coverslip surface by incubating 100 μl of a 1:1000 dilution in PBS (LB30, Sigma Aldrich) for ∼3 minutes. For long measurements (> 3 hours) and long force cycles (non-permissive direction) the tethers were cross-linked to the bottom surface of the flow cell. For this 100 μl of 2% glutaraldehyde solution in PBS was added and incubated for 10 min. After rinsing the flow cell with 0.5 ml of PBS, 50 μl of anti-digoxigenin (50 μg/ml in PBS) was incubated for 30 minutes in the flow cell. The cross-linking reaction with glutaraldehyde was quenched with 1 ml Tris-HCl pH 7.5, as described in (38). The flow cell was then flushed with 1 ml of 10 mM Tris, 1 mM EDTA pH 8.0, 750 mM NaCl, 2 mM sodium azide to remove excess of anti-digoxigenin followed by rinsing with another 0.5 ml of TE buffer (10 mM Tris, 1 mM EDTA pH 8.0, 150 mM NaCl, 2 mM sodium azide). The surface was then passivated with a 10 mg/ml solution of bovine serum albumin (BSA, Fraction V, Carl Roth) in PBS and 5% glycerol for 30 minutes and rinsed with 0.5 ml of TE buffer.

### Flow cell assembly with SpyCatcher surface

For the assembly of flow cells with SpyCatcher surface, we sandwiched a double layer of Parafilm by two #1 coverslips, the top one having one hole at each end serving as inlet and outlet. The flow cell was mounted into a custom-built holder and rinsed with ∼1 ml of PBS. To start the cross-linking reaction, 100 μl of 2% glutaraldehyde solution in PBS was added and incubated for 10 min. The excess was flushed away with 1 ml PBS. 3 μm diameter polystyrene reference beads (LB30, Sigma Aldrich) were attached to the bottom coverslip surface by incubating 100 μl of a 1:1000 dilution in PBS for ∼3 minutes. After rinsing the flow cell with 0.5 ml of PBS, a 100 ug/ml solution of SpyCatcher3 in PBS was added to the flow cell and incubated for 1 hour. The cross-linking reaction was quenched with 1 ml Tris-HCl pH 7.5 (38) and the flow cell was passivated with a 10 mg/ml bovine serum albumin (BSA, Fraction V, Carl Roth) and 10 mg/ml beta-casein (from bovine milk, Sigma-Aldrich) mixture in TE buffer (10 mM Tris, 1 mM EDTA pH 8.0, 150 mM NaCl, 2 mM sodium azide). The mixture was incubated for 30 min in the flow cell and washed away with 1 ml TE buffer.

### Recombinant MTERF1 expression and purification

#### Construction of MTERF1 bacterial expression plasmids

The human MTERF1 gene was obtained from OriGene. The gene was PCR amplified without the mitochondrial signaling sequence and cloned into the p6HIS-SUMO bacterial expression plasmid using BsaI and SalI (39). The final p6HIS-SUMO-MTERF1 construct was confirmed by sequencing by the nucleic acid sequencing facility at PSU.

#### Construction of MTERF1 R162A derivative

A single amino acid substitution was introduced into the MTERF1 coding sequence of p6HIS-SUMO-mTERF1 bacterial expression plasmid using the Agilent QuikChange Site-Directed Mutagenesis Kit, following the manufacturer’s instructions. Mutagenic primers were designed to introduce the desired codon substitution while preserving *E. coli* codon optimization. The following single point mutation was constructed: R162A. The construct was confirmed by sequencing by the nucleic acid sequencing facility at PSU.

#### Expression of human MTERF1

*E. coli* Rosetta (DE3) competent cells were transformed with the p6HIS-SUMO-mTERF1 plasmid (WT and R162A derivative), plated on NZCYM agar plates containing kanamycin (K25, 25 µg/mL), chloramphenicol (C20, 20 µg/mL), and dextrose (0.4%) and grown for 16 h at 30 ^o^C. Rosetta (DE3) cells containing the p6HIS-SUMO-mTERF1 plasmid were grown in 100 mL of media (NZCYM) supplemented with kanamycin (K25) and chloramphenicol (C20) at 37 °C until an OD_600_ of 1.0 was reached. This culture was then used to inoculate 200 mL of K75, C60-supplemented ZYP-5052 auto-induction media at 37 °C (40). The cells were grown at 37 °C to an OD_600_ of 0.8 to 1.0, cooled to 25 °C and then grown for 20 h. After ∼20 h at 25 °C the OD_600_ reached ∼10–15. Induction was verified by SDS-PAGE. Cells were harvested by centrifugation (6000 x *g*, 10 min) and the cell pellet was washed once in 200 mL of TE buffer (10 mM Tris, 1 mM EDTA), centrifuged again, and the cell paste weighed, typical yields were ∼10g per 200 mL culture.

#### Purification of MTERF1

Frozen cell pellets were thawed on ice and suspended in lysis buffer (100 mM potassium phosphate pH 8.0, 500 mM NaCl, 10 mM BME, 20% glycerol) with protease inhibitor cocktail in 5 mL of lysis buffer per 1 gram of cells. The cell suspension was lysed by passing through a French Press at 20,000 psi. After lysis, NP-40 was added to a final concentration of 0.1% (v/v). The lysate was centrifuged at 75,000 x g for 30 min at 4 °C. The clarified lysate was then loaded onto a 5 mL Ni-NTA column at a flow rate of 1 mL/min (approximately 1 mL bed volume per 100 mg total protein) equilibrated with lysis buffer with 5 mM imidazole. After loading, the column was washed with twenty column volumes of lysis buffer with 10 mM imidazole and 5 column volumes of lysis buffer with 50 mM imidazole. The protein was eluted with 5 column volumes of lysis buffer with 500 mM imidazole. Ulp1 (1 µg per mg) was added to the eluted protein to cleave the SUMO fusion and was dialyzed overnight against 1 L buffer A (25 mM HEPES pH 7.5, 500 mM NaCl, 10 mM BME, 20% glycerol) using a dialysis membrane with a 6-8,000 Da MWCO. After dialysis the dialyzed sample was passed through a 1 mL Ni-NTA column to remove the HIS-SUMO protein. The pass-through was collected and fractions containing MTERF1 protein were pooled based on purity (SDS-PAGE) and the protein concentration was determined by measuring the absorbance at 280 nm by using a Nanodrop spectrophotometer using the calculated molar extinction coefficient. Purified protein was aliquoted and frozen at -80 °C until use.

### Oligonucleotide used in the bulk assays

RNA oligonucleotides were from Horizon Discovery Ltd. (Dharmacon). DNA oligonucleotides were from IDT. T4 polynucleotide kinase was from ThermoFisher. [γ-^32^P]ATP (6,000 Ci/mmol) was from Revvity. Nucleoside 5’-triphosphates (ultrapure solutions) were from Cytiva. All other reagents were of the highest grade available from MilliporeSigma, VWR, or Fisher Scientific.

### Annealing of dsDNA substrates

dsDNA substrates (15-bp random double-stranded sequence) were produced by annealing 1 μM DNA oligonucleotides in 25 mM HEPES pH 7.5 and 50 mM NaCl in a Progene Thermocycler (Techne). Annealing reaction mixtures were heated to 90 °C for 1 min and slowly cooled (5 °C/min) to 10 °C.

### Fluorescence polarization experiments on dsDNA binding

Increasing concentrations of MTERF1 were mixed with 1 nM of duplexed FAM-labeled dsDNA in a binding buffer containing 10 mM HEPES pH 7.0, 10 mM BME, 50 mM NaCl, and 0.1 mM EDTA (60 µL volume). Reactions were incubated at room temperature for 1 minute in a black 96-well non-binding plate and then read with a dual wavelength Ex: 485/20 Em: 528/20 fluorescence polarization cube using a Biotek Synergy plate reader. Data from protein titration experiments were fitted with a hyperbola:

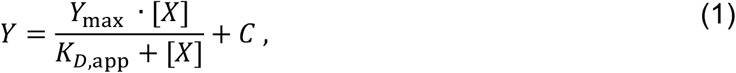

Where X is the concentration of protein, Y is degree of polarization, *K*_*D,app*_ is the apparent dissociation constant, and *Y*_*max*_ is the maximum value of Y, and C is an offset.

### Preparation of dual promoter template

Dual promoter dsDNA templates were prepared by PCR as described previously (41). Templates contained the LSP, HSP, interprotomer region and 22-bp MTERF1 binding site 35 nucleotides from the LSP. PCR products were purified with Wizard SV gel and PCR Clean-up System (Promega), DNA was eluted in 80 µL TE (10 mM Tris-HCl, pH 8.0 buffer and 0.1 mM EDTA) and finally diluted to 1 µM in TE buffer. Extinction coefficients for the DNA construct were calculated with IDT DNA technologies tool (http://biophysics.idtdna.com/UVSpectrum.html

### In vitro transcription assays

Reactions were performed in 1X reaction buffer (10 mM HEPES pH 7.5, 100 mM NaCl, 10 mM MgCl_2_, 1 mM TCEP and 0.1 µg/µL BSA) with 10 μM ^32^P-end-labeled RNA primer (pAAA), 500 μM NTP, and 100 nM DNA template. Reactions were performed by incubating template DNA in reaction buffer at 32 °C for 5 min and then adding in the following order: TFAM (0.5 µM), TFB2M (1 µM), MTERF1 (0 to 10 µM), and POLRMT (1 µM). Between each addition of protein to the reaction there was an incubation time of 1 min. After addition of POLRMT, the reaction was allowed to incubate at 32 °C for 5 to 60 min. At each time point 4 μL of the reaction mix were quenched into 8 μL of stop buffer (79.2% formamide, 0.025% bromophenol blue, 0.025% xylene cyanol and 50 mM EDTA final). Products were resolved by denaturing 20% (37:3, acrylamide:bis-acrylamide ratio) PAGE. Proteins were diluted immediately prior to use in 10 mM HEPES, pH 7.5, 1 mM TCEP, and 20% glycerol. The volume of protein added to any reaction was always less than or equal to one-tenth the total volume. Gels were visualized by using a PhosphorImager (GE) and quantified by using ImageQuant TL software.

### Single-molecule experiments on MTERF1 blocking DNA unwinding

20 μl of streptavidin coated Dynal Dynabeads M270 streptavidin coated magnetic beads (ThermoFisher) was mixed with ∼0.1 ng of DNA (total volume 40 μl) and incubated for ∼5 minutes before rinsing with ∼2 ml of TE buffer to remove any unbound DNA and the magnetic beads in excess. DNA hairpins were tested by looking for the characteristic extension of the correct length (∼1.4 µm at 40 pN) due to the stretching during a force ramp experiment (34). 100 μl mix of 50 nM MTERF1 in reaction buffer (10 mM HEPES pH 7.5, 10 mM MgAc2, 50 mM KGlu, 1g/l BSA and 2 mM sodium azide) was flushed into the flow cell. To assess the strength of the MTERF1-DNA lock formation, we performed several force cycles (> 100) on the same DNA hairpins: 90 s at low force (∼5 pN) to incubate MTERF1, 240 s at a specific force below the critical force of the hairpin to extend it without direct opening and 10 s at ∼35 pN to open all remaining hairpins. The measurement is performed at an acquisition frequency of 58 Hz and shutter time of 17 ms and a constant temperature of 25 °C. A custom written LabVIEW routine controlled the data acquisition and the (x-, y-, z-) positions tracking of both the magnetic and reference beads in real-time (42). Mechanical drift correction was performed by subtracting the reference bead position from the magnetic bead positions and further corrected using an autofocus (along the z-axis) protocol previously described in (43).

### The overdamped harmonic oscillator

The tethered magnetic bead under force can be treated as an overdamped harmonic oscillator experiencing thermal noise in the x- and y-direction (15). For the long pendulum axis, the characteristic timescale of the fluctuations is expressed as

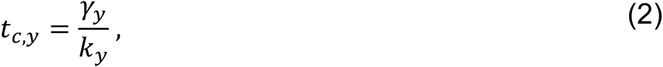

Where *γ*_*y*_ is the drag coefficient along the y-axis and *k*_*y*_ is the trap stiffness of the magnetic field. As the bead can freely rotate on the y-axis (15, 20), the overdamped harmonic oscillator has the combined length of the tether extension *L*_*ext*_ and the radius of the bead *R*, i.e. *L*_*ext*_ *+ R* (20). The motion of the tethered bead originates from the thermal noise and therefore the average kinetic energy of the tethered bead along the y-axis is 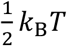, where *k*_*B*_ is the Boltzmann constant and *T* is the temperature. This energy is dissipated in the force field with potential energy 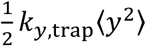, where *k*_*y,trap*_ is the stiffness of the magnetic trap and ⟨*y*^*2*^⟩ is the variance in bead positions along the y-axis. From the conservation of energy, we infer for the trap stiffness

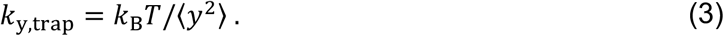

The tether and the bead can be considered as two end-connected rigid pendulums with length *L*_*ext*_ and *R*, and a force *F* exerted along the pendulums in the y,z-plane (**Fig. 3A**). Therefore, the y-components of the pendulums can be respectively expressed as

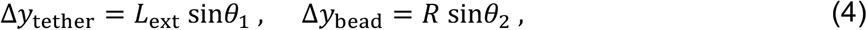

Where *θ*_*1*_ and *θ*_*2*_ are the angles of the tether and bead with respect to the z-axis (**Fig. 3A**). The restoring force on the tether and the bead along the y-axis can be written in terms of the magnetic trap force *F*

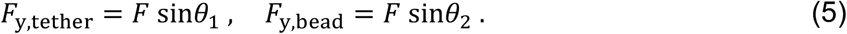

When we combine **Equation 4** and **Equation 5**, we obtain that the spring constants of the oscillators in the y-direction are equal to *F/L*_*ext*_ and *F/R* for the tether and bead, respectively. As the oscillators are connected in series, the combined oscillator has the effective spring constant 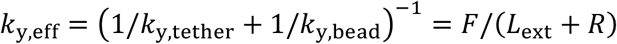. Combining this expression with **Equation 3**, we obtain the expression for the force along the long pendulum axis (44)

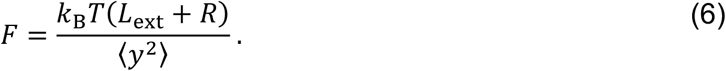

For the characteristic timescale of every overdamped harmonic oscillator holds 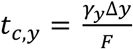. This can be obtained from **Equation 2** and the spring force on the oscillator *F = k* Δ*y, F y* where Δ*y* is the total length of the oscillator, i.e. the length of the double pendulum *L*_*ext*_ *+ R* in the case of tethered magnetic beads along the long pendulum axis (**Results**). Since *γ*_*y*_ and Δ*y* are in principle constant for a constant force *F*, we expect that *t*_*c,y*_ ∝ *1/F* and *log (t*_*c,y*_) ∝ *™log(F)*.

For beads on a long tether, the bead is governed by the Stokes’ drag and the drag coefficient is simply *6πηR*, since the tethers drag is negligible compared to the beads. For short tethers, the bead is close to the surface and a correction factor *C*_||_ is required for translational motion parallel to the surface given by Faxén’s correction (45)

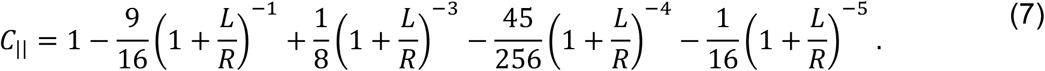

### Force calibration of magnetic tweezers

To measure precisely *L*_*ext*_ and ⟨*y*^*2*^⟩ to calibrate the force from **Equation 6**, the mechanical drift was corrected by subtracting the position of a surface-attached reference bead to the position of the tethered bead, and we used an autofocus during the experiment to further correct the beads position measurements along the z-axis (43). An accurate estimation of ⟨*y*^*2*^⟩ requires a camera shutter time at least 3-fold smaller than the magnetic bead characteristic time *t*_*c,y*_*(F)* to prevent image-blurring (**Fig. S4**) (17). To obtain accurate estimates of the variance on the long pendulum axis ⟨*y*^*2*^⟩ from high to low forces (50-3 pN), we kept the shutter time at 0.4 ms and we measured more than 15,000 frames per magnets height at an acquisition rate of 58 Hz to reduce the statistical error below 3%. As the M270 magnetic beads used in this study are highly monodisperse in size (46), we treated their radius as a constant, i.e. *R = 1*.*4 μ*m. The temperature was kept at 25 °C during the measurement as described in (37) and we used *k*_*B*_*T ≈ 4*.*11* · *10*^*–3*^ pN·*μ*m. The force measurements were performed with the long pendulum axis aligned with the short axis of the flow cell (50 × 5 mm), as noise coupling was minimal along this axis.

### Data processing

The activity traces were first corrected from the mechanical drift by subtracting the reference bead position to the tethers position. The median of the high force (35 pN) and low force (5 pN) parts of the z-traces were subsequently obtained, to determine the midpoint of the z-traces. From the parts of the z-traces at the intermediate forces, regions of 0.2 μm around the midpoint were extracted to determine the dwell-time at the target region, where the DNA hairpin was opened up to the binding site of MTERF1. The dwell-times of all traces for a given experimental condition were assembled into a single distribution and further analyzed using a maximum likelihood estimation (MLE) fitting routine to extract the parameters from the dwell-time fit-function.

### Maximum likelihood estimation fitting routine

The dwell-time distributions were fitted to the experimentally collected dwell-times *{t*_*i*_*}* by maximizing the log-likelihood function

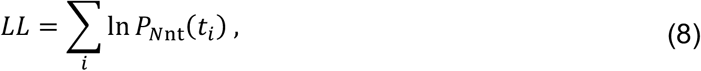

Where *P*_*Nnt*_ represents the probability of every dwell-time *t*_*i*_ in the empirical distribution based on the dwell-time fit-function, We calculated the statistical error on the parameters by applying the MLE fitting procedure on 100 bootstraps of the original data set, and reported the standard deviation for each fitting parameter.

To be able to compare fits of models with different number of parameters, we calculated the Bayesian Information Criterion (BIC)

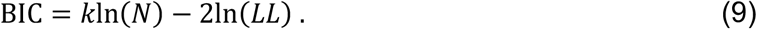

This criterion compares the log-likelihood *LL* to the number of parameters in the model *k* and the number of datapoints fitted *N* to account for the fact that using more parameters can lead to overfitting. The model fit with minimized BIC is considered the best fit with optimal number of parameters for the set of model fits tested (47).

## RESULTS

### MTERF1 efficiently terminates LS-transcribing mtRNAP

To investigate the mechanism of transcription termination by MTERF1, we recombinantly expressed in *E. coli* and purified the human MTERF1 (**MATERIALS AND METHODS**) (**Fig. S1A**). Wild-type MTERF1 showed an apparent equilibrium binding constant to the termination site of *K*_*d,app*_ = (0.33 ± 0.03) μM (**Fig. S1B**), which is in close agreement with a previous report (27). Mutating the binding pocket of T3243 from arginine to alanine (R162A) did not change the binding affinity (**Fig. S1B**), indicating that MTERF1 binding affinity does not originate from the everted bases. We then investigated whether MTERF1 was able to terminate LS-transcribing mtRNAPs, while not affecting HS-transcribing mtRNAPs. We designed a DNA construct including both LSP and HSP, and a termination site downstream the former (**Fig. 1C**) (**MATERIALS AND METHODS**). We performed run-off mtRNAP transcription experiments on this construct using *de novo* transcription initiation conditions (**MATERIALS AND METHODS**), while titrating wild-type (WT) MTERF1 concentration from 0 to 5 µM (**Fig. 1D**). We observed that LS-transcribing mtRNAPs were strongly affected by MTERF1, with nearly complete transcription termination at 2.5 µM of MTERF1 (**Fig. 1DE**). The R162A MTERF1 mutant decreased the transcription termination efficiency by nearly half (**Fig. 1DE**), confirming the importance of the T3243 everted base stacking to terminate mtRNAP (27). Interestingly, we also noticed that the band corresponding to HS-transcribing mtRNAP decreased in intensity when increasing MTERF1 concentration (**Fig. 1D**). This suggests that a significant fraction of LS-transcribing mtRNAP remained trapped on the DNA construct upon collision with MTERF1, indicating that these mtRNAPs were not recycled. Our bulk assay clearly shows that MTERF1 binds with high affinity to the termination site and that T3243 base eversion and stacking is important to terminate transcription, while not affecting MTERF1 binding affinity.

### A magnetic tweezers assay to interrogate MTERF1 directional lock

We then employed a high-throughput magnetic tweezers assay to investigate the mechanism of MTERF1 transcription termination. We used an assay that was previously described to interrogate the Tus-*Ter* polar roadblock origin (8). Specifically, we designed a DNA hairpin encoding a termination site recognized by MTERF1 in either orientation (permissive or non-permissive) in the middle of the hairpin stem (**Fig. 2B**). We employed these hairpins to investigate whether MTERF1 bound to the termination site blocks DNA unwinding directionally (**Fig. 2AB**). To this end, we performed force-jump experiment where we started the force cycle at low force (5 pN) for 90 s, i.e. hairpin closed, to enable MTERF1 binding to the termination site. This was followed by a jump to a higher force that is above the critical value at which the hairpin opens (∼16 pN), enabling hairpin unwinding up to the bound MTERF1 (**Fig. 2A**). We recorded the time for MTERF1 to unlock the high force jump. Finally, the force was increased to 35 pN for 10 s, such that all the hairpins were fully open and all MTERF1 unlocked from the hairpins stem, before restarting the cycle at the low force. This force cycle was repeated many times (> 100) to record as many MTERF1 unlocking events as possible on a single DNA hairpin.

**Figure 2:**
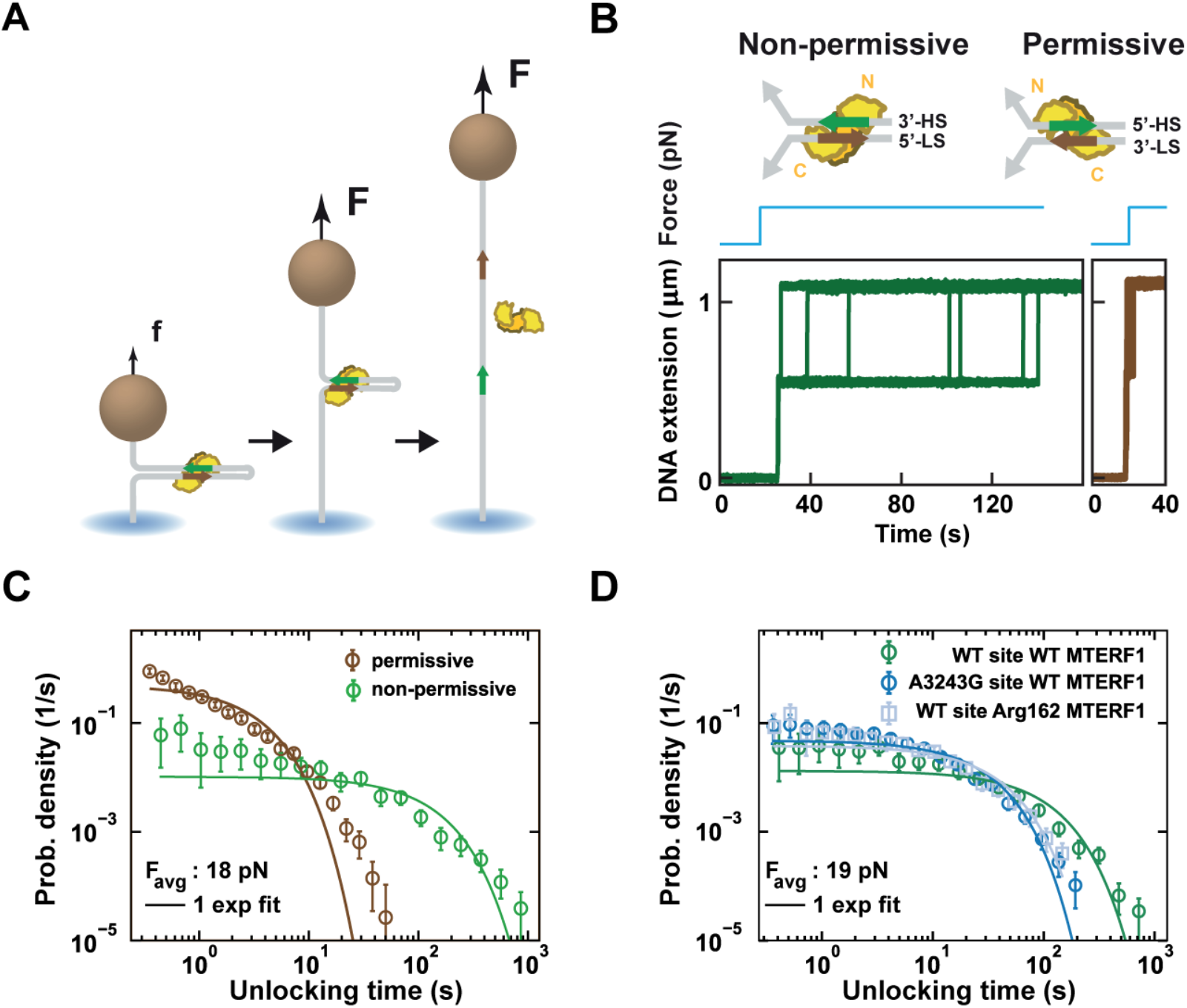
A force-jump magnetic tweezers assay reveals asymmetric stability of MTERF1 association to the termination site as a function of DNA unwinding direction. **(A)** Force-jump hairpin opening assay to apply directional DNA unwinding on MTERF1-termination site complex. The force increases from low (f), where the hairpin is closed and MTERF1 can bind to the termination site, to high (F) in one step, enabling hairpin opening until the DNA fork reach MTERF1. Full hairpin opening occurs when MTERF1 dissociates. The force is then brought back to f and the cycle is repeated. **(B)** Example traces for MTERF1 associated with the termination site with a critical force on the hairpin for the permissive and non-permissive orientation. (Top) Schematic representations of DNA directional unwinding with MTERF1 bound to the termination site, in the non-permissive (left) and permissive orientation (right) with the binding sequences on LS and HS indicated as brown and green arrows, respectively. **(C)** Unlocking time distributions for the permissive (N=3813) and non-permissive (N=363) orientation at a constant average force of *F*_*avg*_ = 18 pN. **(D)** Unlocking time distributions using non-permissive hairpins and average force *F*_*avg*_ = 19 pN, for wild-type (WT) termination site and WT MTERF1 (N=671), with A3243G termination site mutant and WT MTERF1 (N=1133), and WT termination site and R162A MTERF1 mutant (N=667). (C, D) The solid lines are single exponential decay fits to the data (circles of the same colour). The error bars represent the standard error of the mean (SEM) extracted from 1000 bootstraps.

Performing the force cycles described above (**Fig. 2A**), we observed a significant difference in the unlocking times between the hairpins with the termination site in either the permissive or the non-permissive orientation (**Fig. 2B**). In the former case, the hairpin remained half open only for a few seconds, whereas in the latter case, the hairpin remained half open for tens to several hundreds of seconds. At a critical force of 18 pN, MTERF1 barely blocked the hairpin opening in the permissive orientation, with an unlocking rate of *(0*.*43 ± 0*.*01)* s^-1^ (mean ± standard deviation from 1000 bootstraps, **MATERIALS AND METHODS**), while in the non-permissive orientation MTERF1 unlocked at a ∼40 times slower rate of *(0*.*010 ± 0*.*001)* s^-1^ (**Fig. 2C**). From these experiments, we concluded that MTERF1 directionally blocks DNA unwinding at the termination site.

### Mutating MTERF1 binding pocket and associated base destabilizes MTERF1 lock in the non-permissive orientation

We next interrogated how the residues interacting with the everted bases A3243 and T3243 in MTERF1 stabilized the lock. To this end, we employed the R162A MTERF1 mutant that disrupts transcription termination *in vitro* (**Fig. 1DE**) (29). This mutant showed a ∼3-fold faster unlocking compared to the wild-type MTERF1, i.e. *(0*.*037 ± 0*.*002)* s^-1^ vs. *(0*.*013 ± 0*.*001)* s^-1^, respectively (**Fig. 2D**). We next modified the termination site at the everted bases AT3243 (**Fig. 1B**) to CG3243, as this mutation was shown to partially abolish transcription termination (27). Here again, we observed a higher unlocking rate than for the wilt-type sequence, i.e. *(0*.*047 ± 0*.*002)* s^-1^ (**Fig. 2D**). When we combined both the MTERF1 and termination site mutants, we measured an even faster unlocking rate for MTERF1, i.e. *(0*.*149 ± 0*.*020)* s^-1^ (**Fig. S2A**). Mutating a non-everted base in the termination site (GC3249 to AT3249) resulted in a lesser impact on MTERF1 unlocking rate than mutating the everted base, i.e. *(0*.*022 ± 0*.*002)* s^-1^ (**Fig. S2B, Table S2**). Altogether, this data confirms that the stacking interactions of T3243 with R162 are essential to form a stable lock on the termination site.

### *In-situ* force calibration for accurate force spectroscopy with magnetic tweezers

We next set out to determine the energy landscape of MTERF1 unlocking from the termination site in the non-permissive orientation when the hairpin experiences a constant force. As expected for such experiment (48), we found that the unlocking rate is highly force dependent (**Fig. S3A**). Surprisingly, the unlocking dwell time were not described by a single exponential distribution, as expected for a reaction with a single rate-limiting transition, but rather required two or more exponentials to be accurately represented (**Fig. 2C, Fig. S3A**). As the distributions were assembled by combining all the dwell times from every tether, with different magnetic beads, we asked ourselves whether the multi-exponential distribution resulted from the bead-to-bead difference in force. Indeed, we previously showed that the dispersion in force between the beads is ∼10% standard deviation relative to the mean (17), for a given distance of the magnets to the beads. This force dispersion may be sufficient to observe multi-exponential behavior. To demonstrate this, we simulated three kinetic processes that are single exponentially distributed with different rates (mimicking the MTERF1-DNA unlocking events on three beads experiencing forces with 10% difference between the beads, i.e. 18, 20 and 22 pN). We obtained a multi-exponential distribution for the MTERF1 unlocking times when combining the three distributions (**Fig. S3BC**). To overcome this issue, we set out to establish an *in-situ* force calibration method for each tether in the field of view instead of using an average force calibration standard, as usually done (8).

The motion of the tethered magnetic bead experiencing an attractive force from the magnets can be described by an inverted pendulum (19). The magnetic field direction is defined by the orientation of the two magnets (**Fig. 3A**), which results in the magnetic bead being clamped in the direction parallel to the magnetic field (17). In the direction perpendicular to the field lines the bead is free to rotate around the magnetic field line (y-axis in **Fig. 3A**), therefore defining a short and a long pendulum axis, respectively. While force calibration in magnetic tweezers is usually treated along the short pendulum axis (15–18), the long pendulum axis provides a more accurate force calibration for short tethers (20). The equipartition theorem provides a mathematical description of the force *F* as a function of the tether extension *L*_*ext*_, the magnetic bead position variance along the long pendulum axis ⟨*y*^*2*^⟩, the magnetic bead radius *R* and the thermal energy *k*_*B*_*T* (**MATERIALS AND METHODS**)

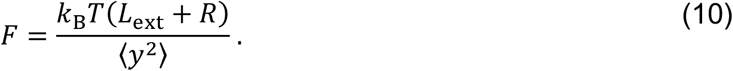

**Figure 3:**
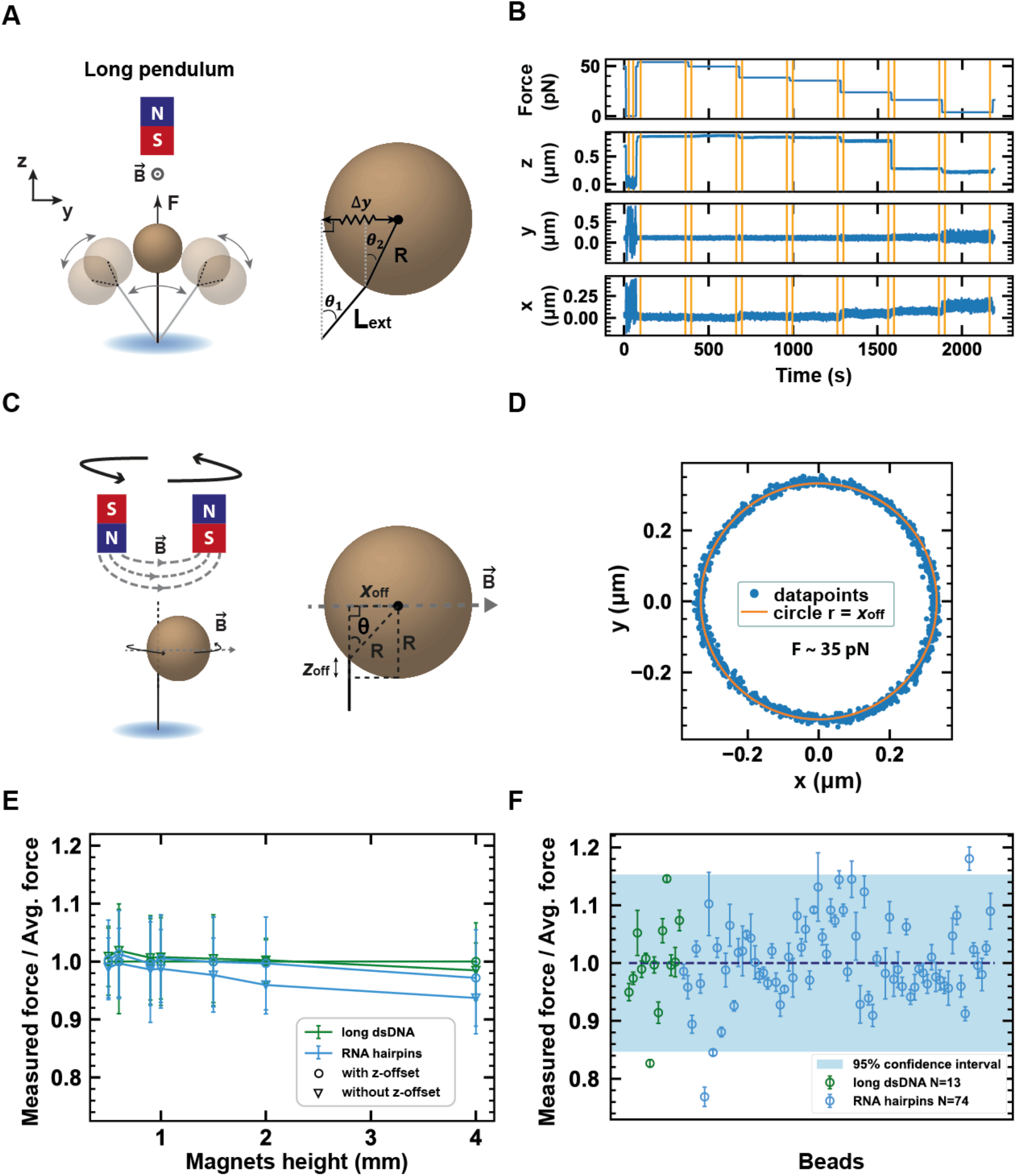
In-situ force calibration enables accurate force spectroscopy measurement for high-throughput magnetic tweezers. **(A)** Left: schematic of the long pendulum description. Right: representation of the tether side attachment for the long pendulum axis (y-axis). **(B)** Example time traces of the x, y and z position at the indicated expected forces. Each force is measured for 5 minutes. The jump in the z-axis trace at ∼1600 s indicates hairpin closing. **(C)** Schematic of the slow rotation measurement (left). Geometric description of the magnetic bead side attachment position to calculate the bead height correction *z*_*off*_ (right). **(D)** Slow tethered magnetic bead rotation extracted while rotating the magnets from -10 to +10 turns at 0.1 turn/s and acquiring images at 58 Hz (blue dots). The rotation is performed under ∼35 pN force applied to the tether. The orange solid line is a fit to the data to measure the radius *x*_*off*_. **(E)** Forces estimated for different magnets heights from measurements on the long pendulum axis with or without *z*_*off*_ correction (circles or triangles, respectively) on long dsDNA (green) and short RNA hairpins (blue) normalised to the average force with *z*_*off*_ on the long dsDNA. **(F)** The standard deviation in force divided by the mean (relative standard deviation) estimated per bead attached to either the long dsDNA tethers or the short RNA hairpins per magnet height. The blue shade represents the 95% confidence interval for the average force calibration of all the beads.

*L*_*ext*_ is obtained from subtracting the lowest z-position of the magnetic bead at zero force to the median z-position of the magnetic bead at a given magnets height and force *F* (**Fig. 3B**), i.e. *L*_*ext*_(*F*) = *z*(*F*) − *z*(0). Using **Equation 10** directly, we observed that the force measured using either the short or long pendulum description is identical when using long tethers (∼7 µm contour length **Fig. 3E**). However, for short RNA hairpin tethers (∼0.5 µm contour length), we noticed a consistent underestimation of the force (**Fig. 3E**). This discrepancy originates from the tether being side attached to the bead, and not to the south pole, biasing the tether extension measurement (49) (**Fig. 3C**). This is specifically important when *L*_*ext*_ is comparable to or smaller than *R* (**Equation 10**), which is the case for the hairpins (**Fig. 3B**). Shon and colleagues (50) showed that the distance from the bead south pole to the attachment point of the DNA along the z-axis *z*_*off*_ can be described by

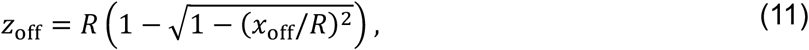

Where *x*_*off*_ is the radius of the circle described by the magnetic bead when slowly rotating the magnets (**Fig. 3CD**) (50). Few beads on the edge of the field of view showed other patterns than a circle, e.g. Pascal limaçon (51), indicating that the beads are being pulled in a direction not strictly parallel to the z-axis, and were discarded from further analysis. The values of *x*_*off*_ at each force *F* were obtained from the reference value obtained by slow rotation at 35 pN, i.e. *x*_*off*_(35 pN), and the median x-position during the force measurement, i.e. *x*(*F*), as

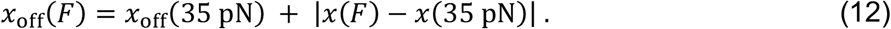

Subsequently, *z*_*off*_ is obtained by substituting **Equation 12** into **Equation 11** and the true tether extension can now be expressed as

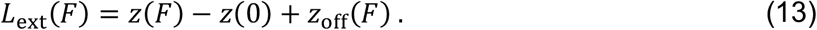

Using the corrected value of *L*_*ext*_ into **Equation 10**, the force calibration when using short RNA hairpins matched the calibration when using long dsDNA tethers (**Fig. 3E**).

To further interrogate whether our approach can be used for accurate force estimation, we evaluated the characteristic timescale for the tethered bead along the long pendulum axis as a function of the measured force (**MATERIALS AND METHODS**)

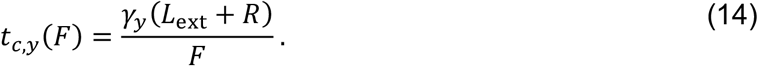

From the expression in **Equation 14** we obtain that for tethers with *L*_*ext*_ *<< R*, the characteristic timescale along the long pendulum axis is dominated by the magnetic bead radius. Force calibration using this method can therefore accommodate longer shutter time than the short pendulum without affecting the measurement (36). Looking at the trends in the characteristic timescales, we noticed that the slope in *log* (*t*_*c,y*_(*F*)) vs. *log*(*F*) is close to -1 for the long DNA tethers (**Fig S4A**), as expected for an overdamped harmonic oscillator (**MATERIALS AND METHODS**). However, for closed hairpins, we observed a clear drop in the calculated characteristic timescale relative to the measured force (**Fig. S4B**). For such short tethers, *γ*_*y*_ must be corrected for the additional friction induced by the magnetic bead motion parallel to a nearby surface of the flow cell (45). This is described by the Faxén’s law and results in a correction factor *C*_||_ on *γ*_*y*_ (**Equation 7, MATERIAL AND METHODS**)

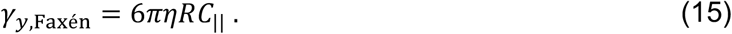

Using *γ*_*y*,Faxén_, *log* (*t*_*c,y*_(*F*)) values converged to a linearly decreasing trend with *log*(*F*) with slope -1 as for the long DNA tethers (**Fig. S4B**). Our description of the magnetic tweezers along the long pendulum axis shows that the system bead—short tether can be described as a single harmonic oscillator (**MATERIALS AND METHODS**). This interpretation could also be applied to other force spectroscopy techniques described by an inverted pendulum, e.g. acoustic force spectroscopy (2). In conclusion, the forces are correctly calibrated with our present method (**Fig. 3**).

Next, we benchmarked the precision of the *in-situ* force calibration. To this end, we calibrated the force for each tethered magnetic bead in the field of view. We found that the relative standard deviation to the mean force (relative error) decreases with the statistical error as 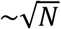 over the range measured (**Fig. S5**). For the calibration standard, however, the force is obtained by combining the calibration from multiple beads, the relative error saturates at a value of ∼0.08, i.e. ∼8% standard deviation compared to the mean (**Fig. S5**). Performing this *in-situ* force calibration on long dsDNA tethers (20.6 kbp, 7 μm contour length) and short RNA hairpins (∼1.6 kbp handles, ∼0.5 and ∼1 μm extension when either closed or open, respectively) (**MATERIALS AND METHODS**), we obtained very similar force estimates from low to high magnet heights using the long pendulum theory (**Fig. 3E**). Furthermore, the relative error obtained with *in-situ* force calibration is 1-3%, while the error was ∼8% for the calibration standard, since we overcome the bead-to-bead variation (**Fig. 3F**). In conclusion, the *in-situ* force calibration strategy we describe here provides a simple and robust strategy to perform high-precision force measurements that are only limited by the statistical error.

### SpyTag–SpyCatcher surface attachment strategy increases the tethers lifetime

To investigate whether MTERF1 unlocking in the non-permissive direction is described by a single exponential distribution, the hairpins experience dozens of cycles lasting 240 s at a critical force (15-20 pN). This prolonged exposure to high force results in digoxygenin–anti-digoxigenin bonds to eventually break before enough statistics is acquired for each hairpin (**Fig. S6**) (38). We therefore had to increase the tether lifetime to collect MTERF1 unlocking times through force-jump cycles. To this end, we employed a recently developed glutaraldehyde cross-linking method (38), which we adapted by replacing anti-digoxigenin by SpyCatcher and digoxenin by SpyTag, providing a covalent bound between the tether and the nitrocellulose cross-linked SpyCatcher (**MATERIALS AND METHODS**). The other handle was biotin-modified to attach the streptavidin coated magnetic bead (**MATERIALS AND METHODS**). The SpyTag-SpyCatcher reaction is specific and limited by diffusion (32, 33), resulting in a very high yield of tethers bound to the flow cell surface. This strategy offered a stable tether attachment, with the average tether lifetime increasing from (15418 ± 1145) s to (25828 ± 3365) s for repeated force-jump experiments, with tethers lasting more than 11 hours (**Fig. S6**).

### Probing the energy landscape of the MTERF1 lock

Combining accurate, *in-situ* force calibration and increased tether lifetime, we were able to investigate the energy landscape of MTERF1 unlocking in the non-permissive orientation. To this end, we started the experiment with *in-situ* force calibration, followed by dozens of force-jump cycles using the hairpin encoding the termination site in the non-permissive orientation and in the presence of MTERF1 in the reaction buffer. We obtained the mean and standard error of the mean (SEM) for the unlocking rate *k*_unlock_ as a function of the *in-situ* calibrated force for each hairpin by calculating the inverse average unlocking time from 100 resampled unlocking time distributions (**Fig. 4A, MATERIALS AND METHODS**). In a log-linear representation, the data were well-fitted by a linear regression representing the Arrhenius equation for a single rate

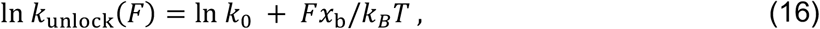

**Figure 4:**
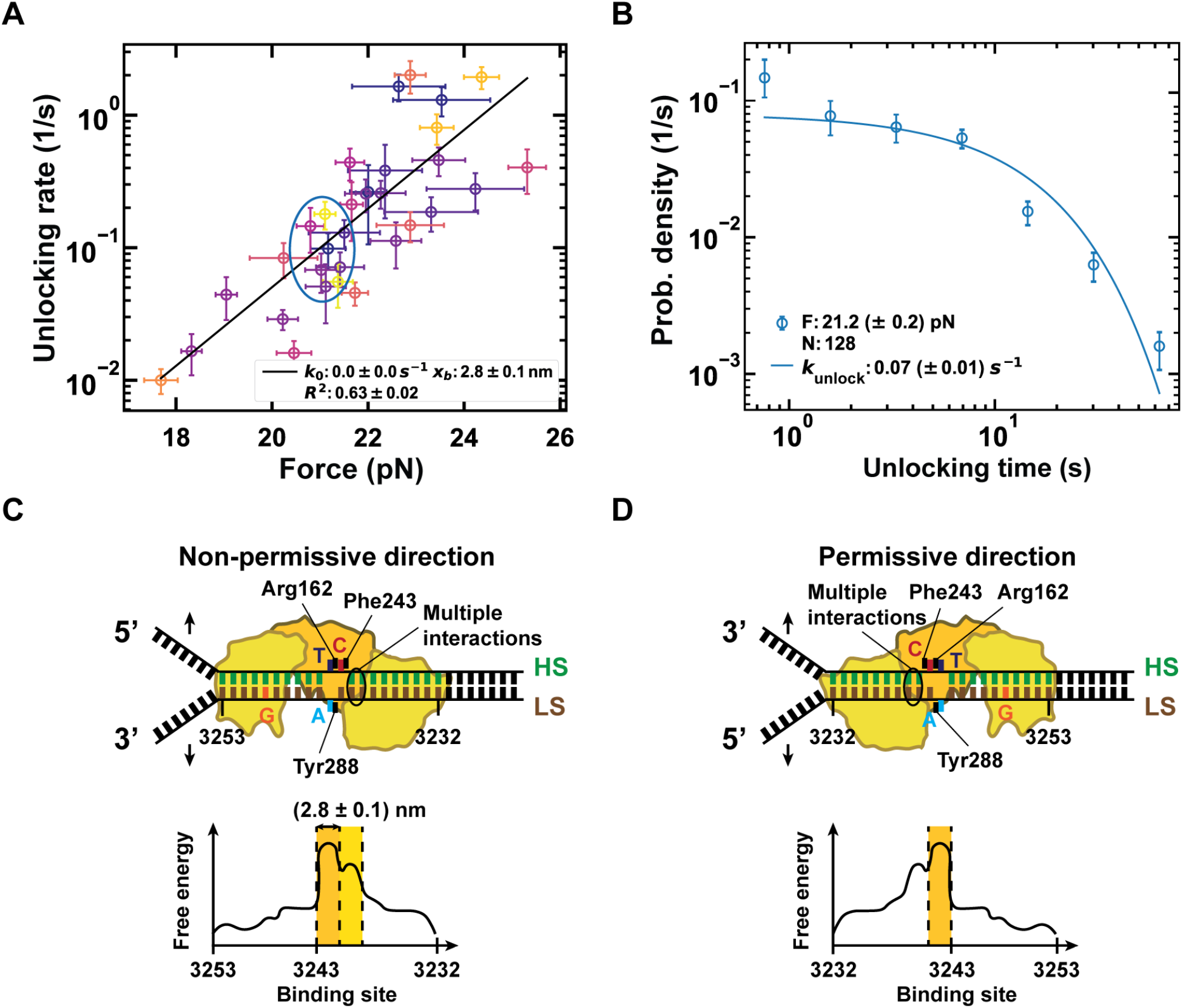
Asymmetric interactions between MTERF1 and the termination site explains MTERF1 polar roadblock activity. **(A)** The unlocking rates as a function of the *in situ* measured force on individual tethers. Each circles represents a single DNA hairpin tethered magnetic beads. The solid line is linear regression of the Arrhenius law to the data that reports a distance to the transition state of (2.8 ± 0.1) nm as fit parameter. The vertical error bars are one standard error of the mean (SEM) and the horizontal bars are one standard deviation from the mean in force. The blue region shows the selection of beads used for (B). **(B)** The combined unlocking time distribution from the beads in the blue region from (A). The solid line is a single exponential probability density function fitted to the data with rate *k*_*unlock*_. **(C, D)** Schematics of the interactions of MTERF1 with the termination site indicating the everted bases (C3242 in red, A3243 in light blue and T3243 in dark blue) and the multiple weaker interactions for the non-permissive (C) and permissive (D) orientation. (Bottom) Free-energy landscapes for the DNA unwinding in either the non-permissive (C) or permissive (D) direction. The distance to the transition state is indicated over the region of the strong interaction in orange, i.e. base stacking, and the region of the weaker interactions is indicated in yellow.

Where *k*_0_ is the rate at zero force and *x*_*b*_ is the distance to the transition state. We extracted *k*_*0*_ = (0.0 ± 0.0) s^-1^, indicating the MTERF1 lock is very stable at *F* = 0 pN, and *x*_*b*_ = (2.8 ± 0.1) nm, which is comparable to the distance separating the three everted bases in the termination site when applying ∼20 pN force (**Fig. 1B**).

For a single transition, the probability density function (pdf) of the unlocking time *t* at a given force *F* is described by a single exponential such as

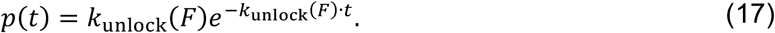

By combining the unlocking times for different hairpin-tethered magnetic beads, for which we estimated the force to be *F = (21*.*2 ± 0*.*2)* pN (**Fig. 4A**), we showed that the resulting distribution is indeed well-described by a single exponential pdf (**Equation 17, Fig. 4B**). This shows that MTERF1 unlocking in the non-permissive direction is dominated by a single transition and highlights the precision of our *in-situ* force calibration method.

## DISCUSSION

In the present study, we showed that MTERF1 terminates transcription and locks on the termination site to halt DNA unwinding in a directional manner. The average unlocking time in the non-permissive direction was ∼40 times longer than in the permissive direction at 18 pN unwinding force (**Fig. 2AB**), showing that MTERF1 is a polar roadblock. We further showed that removing the interactions between MTERF1 and the everted bases in the termination site, by either mutating the bases or the relevant residues in MTERF1 binding pockets, reduced transcription termination efficiency *in vitro* (**Fig. 1CDE**) and destabilized the lock (**Fig. 2C**). The destabilization was even more potentiated when both the sequence and the binding pocket were mutated (**Fig. S1A**), supporting previous ensemble studies describing that base eversion and stacking is essential for stable MTERF1 binding (27). To further explore the energy landscape of MTERF1 locking on the termination site, we established two methods for force spectroscopy experiments with magnetic tweezers. We first developed an *in-situ* force calibration to precisely measure the force experienced by each individual tether in the field of view, even when using short tethers, with an accuracy only limited by the statistical error (**Fig. 3, Fig S4**). Subsequently, we implemented a covalent DNA tether–glass coverslip attachment strategy using a SpyTag–SpyCatcher covalent reaction that enables force spectroscopy measurements for more than 11 hours (**Fig. S5**). These two methods empowered us to perform precise force spectroscopy measurements on the MTERF1 unlocking time using high-throughput magnetic tweezers and show that MTERF1 unlocking is well-described by a single transition (**Fig. 4AB**).

MTERF1 bound to the termination site efficiently terminates mitochondrial LS transcription, but does not hinder HS transcription (24, 25), and similarly pauses the mitochondrial replisome by halting TWINKLE helicase (30, 31). The mechanism behind this polar roadblock is therefore unlikely to be related to specific protein-protein interactions. Therefore, the interactions between MTERF1 and the termination site are responsible for such directional transcription termination or replication pausing. A structural study has shown that MTERF1 forms multiple weak interactions with the DNA bases next to the everted bases on the LS 5’-end of the binding site (**Fig. 4C**), whereas lesser interactions are located on the LS 3’-end (27). While the everted nucleotides contribute the most to the lock stability (**Fig. 1B**), multiple weak interactions can synergize with the everted bases to either stabilize or weaken the lock, depending on the DNA unwinding direction. We propose that an asymmetry in the number of weak interactions flanking strong interactions resulting from the everted bases either potentiate or weaken the lock (**Fig. 4C**). On the one hand, the weak interactions increase the stability of the lock in the non-permissive orientation, while, on the other hand, they provide a progressive increase in the barrier, favoring rapid unlocking (**Fig. 4D**). Future work monitoring real-time collisions between a translocating mtRNAP and MTERF1 at high spatiotemporal resolution will further inform on the exact mechanism of transcription termination. We propose that our model is applicable to other polar roadblocks reported in the literature. The Tus-*Ter* lock that terminates transcription in bacteria, for example, also shows a high density of weak interactions present immediately after the single everted base in the non-permissive direction (8, 52). Furthermore, CRISPR-Cas9 forms a directional roadblock when unzipping DNA, and Zhang and colleagues found indications that weak interactions downstream of the PAM site are responsible for Cas9 polar roadblock activity (53). Future work will elucidate whether other directional roadblocks share a similar mechanism, such as the directional transcription terminator factor-I (TTF-I) that terminates RNA polymerase I transcription in higher eukaryotes (54, 55).

Large data sets with precise force measurement are required to statistically model mechanochemical reactions in their full complexity using a data-driven approach. High-throughput magnetic tweezers are particularly well-suited for such measurements, but they often rely on force calibration standards obtained a priori from averaging the force calibration obtained from many beads using long DNA tethers (7, 8, 11, 16–18). This approach limits the force precision to ∼10% relative error due to the bead-to-bead magnetic content difference (17). While a power-spectrum approach for force calibration has been derived for short tethers by Seidel and colleagues (20), such a method may be difficult to implement for non-physicists and therefore our simple – yet precise – force calibration methodology will be of broad interest. Our approach will enable accurate reconstructions of complex reaction energy landscape with high confidence.

We also provide a strategy for long-lived DNA tether attachment to the glass surface. Several articles have described covalent, long-lived binding strategy for surface-tether attachment for magnetic tweezers experiments (11, 56, 57). However, they often suffer from low tether density due to the low yield of covalent bond reactions. By adapting the SpyTag– SpyCatcher attachment strategy, already used in protein force spectroscopy (33), we benefitted from very efficient binding only limited by diffusion (32, 58). This strategy enables strong covalent bond formation and high tether attachment yield, while it only requires the addition of a SpyTag at the 5’-end of a PCR primer used during the DNA handle fabrication. Despite the improvement in tether lifetime, they still detached from the surface, as crosslinking does not perform as well as directly linking the tethers to the glass surface (38, 57). Future work will adapt the SpyTag–SpyCatcher strategy on a, e.g. pegylated surface, to obtain a covalent linkage to the glass surface while maintaining the reaction yield (56).

Our present work will impact the single-molecule force spectroscopy field and the mitochondrial transcription community. We anticipate that our technical advances will be broadly adopted by the magnetic tweezers force spectroscopy community and will facilitate the investigation of other polar roadblocks, such as TTF-1.

## Supporting information

Supplementary Informations

## RESOURCE AVAILABILITY

### Data statement

Further information and requests for resources should be directed to and will be fulfilled by the lead contact, Dr. David Dulin.

### Data and code availability

This manuscript is supplemented with three data analysis programs in Python as discussed in the main text. The data and analysis scripts used for every figure and table in this manuscript will be made available as soon as the manuscript will be accepted.

## ACKNOWLEDGMENTS

We would like to thank Mohammed Sadegh Feiz for reviewing the scheme for the orientation of the magnetic field. DD was supported by the Interdisciplinary Center for Clinical Research (IZKF) at the University Hospital of the University of Erlangen-Nuremberg, BaSyC – Building a Synthetic Cell” Gravitation grant (024.003.019) of the Netherlands Ministry of Education, Culture and Science (OCW) and the Dutch Organization for Scientific Research (NWO). DD was supported by German Research Foundation (DFG) DU1872/3-1.

## AUTHOR CONTRIBUTIONS

DD, CEC and JJA designed the research. DD, EO and PPBA designed the single-molecule experiments. FS, SQ and QS made DNA constructs. BJ, JJA and CEC provided the purified proteins. BJ and JJA performed the bulk MTERF1-dsDNA binding assays and the *in vitro* transcription experiments. PPBA designed and implemented the *in-situ* force calibration method and DB performed initial experiments. PPBA and EO acquired and analyzed the force-jump experiments with MTERF1. PPBA and DD wrote the article.

## DECLARATION OF GENERATIVE AI AND AI-ASSISTED TECHNOLOGIES

During the preparation of this work no AI-assisted technologies have been used.

