## Supplementary Informations for "Accurate single-bead force calibration in high-throughput magnetic tweezers reveals the mechanism of directional transcription termination by MTERF1"

**This document includes:**

Figures S1 to S6

Tables S1 and S2

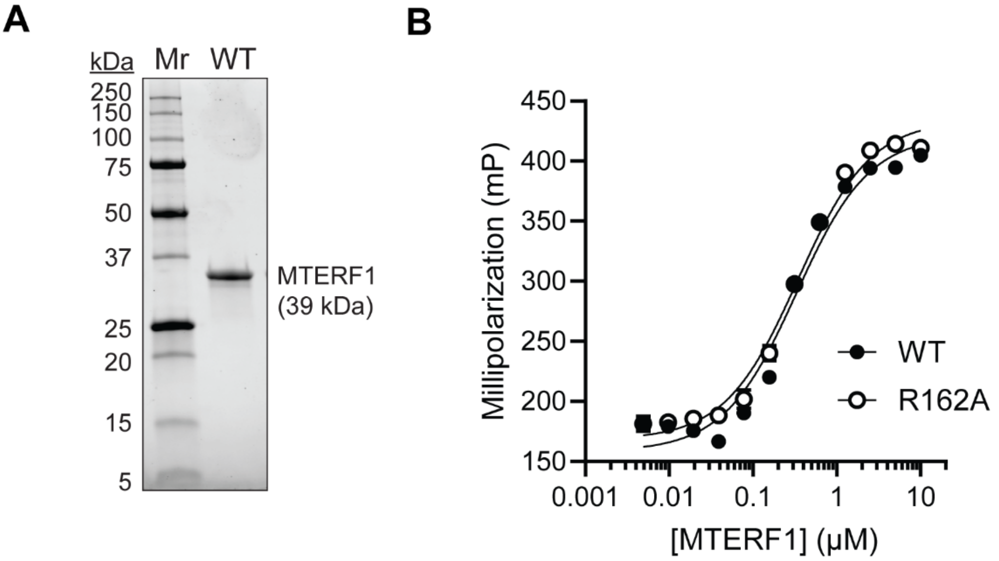


**Figure S1: MTERF1 purification and activity tests. (A)** SDS-PAGE of purified human MTERF1 recombinantly expressed in *E. Coli*. 2.5 μg of purified mTERF1 (WT) is shown. Molecular weight markers (Mr) and corresponding molecular weights are indicated. **(B)** Fluorescence polarization (mP) for increasing concentrations of MTERF1 (WT and R162A) bound to 1 nM FAM–labeled duplex DNA. Fluorescence polarization values (mP) were fitted with **Equation 1** to obtain apparent dissociation constants ($K_{D,\mathrm{app}}$). For WT, $K_{D,\mathrm{app}}$ = (0.33 ± 0.03) μM, while for R162A mutant, $K_{D,\mathrm{app}}$ = (0.34 ± 0.02) μM. The circles with error bars represent mean ± standard deviation (*n* = 3).

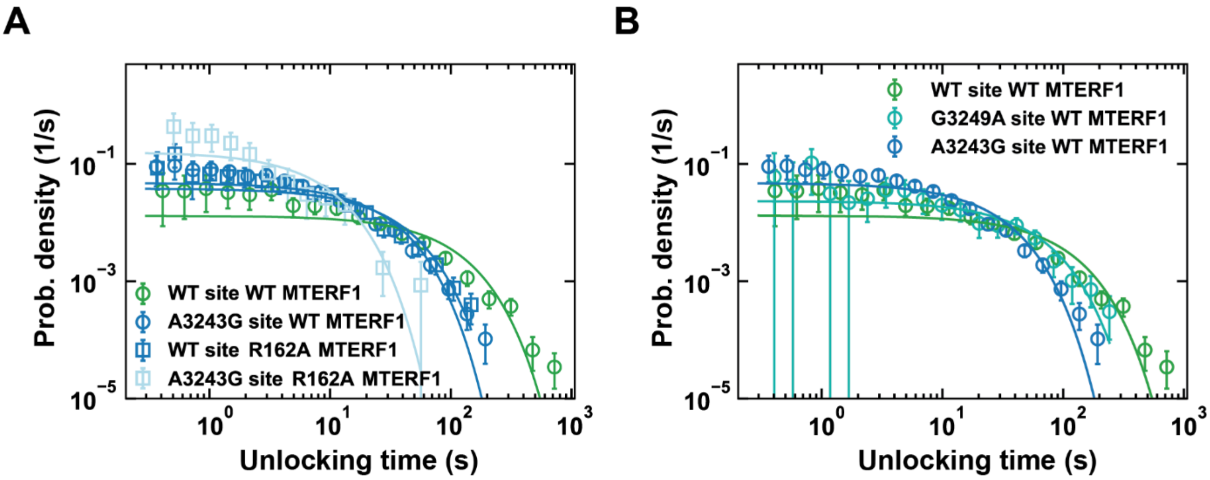


**Figure S2: Unlocking time distributions for different mutations on MTERF1 and the termination site compared to the wild-type protein and sequence. (A)** Distributions for the WT MTERF1 on the WT (green) and A3243G mutated target site compared to the Arg162 mutated (R162A) MTERF1. **(B)** Distributions for the wild-type (WT) MTERF1 on the G3249A and A3243G mutated target site (turquoise and blue) compared to the WT target site (green). The circles with error bars in the distributions represent the mean and SEM from the unlocking times and the lines represent the single exponential probability density function, i.e. the fit curve.

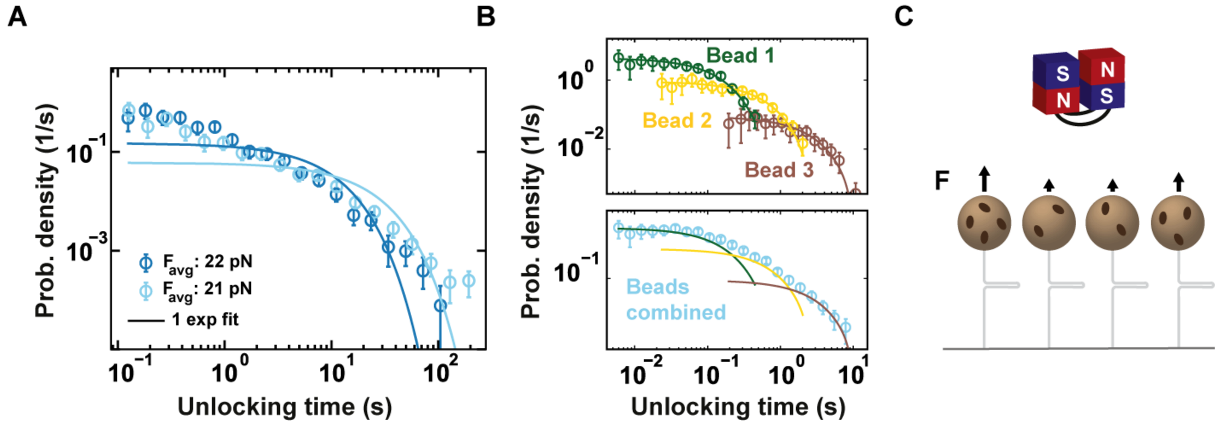


**Figure S3: Combining non-permissive MTERF1 unlocking times for multiple hairpin-tethered magnetic beads results in multi-exponential distributions. (A)** Experimental unlocking time distributions for all beads combined at two magnet heights on the non-permissive hairpin with the corresponding average stretching force indicated (dark and light blue). The solid lines are single exponential fits to the respective distributions. **(B)** (Top) Distributions for simulated unlocking times with three different rates (mimicking unlocking times from three beads experiencing 10% different forces at the same magnet height), assuming unlocking can be described by a single force dependent rate. (Bottom) Unlocking times distribution for the three forces combined. The combination of unlocking events from different beads results in a power law tail in the distribution for long timescales. **(C)** Cartoon of tethered beads stretched under the pulling force from two antiparallel magnets above the flow chamber. The beads experience different forces for the same magnet height since they have different magnetic content (dark brown spots).

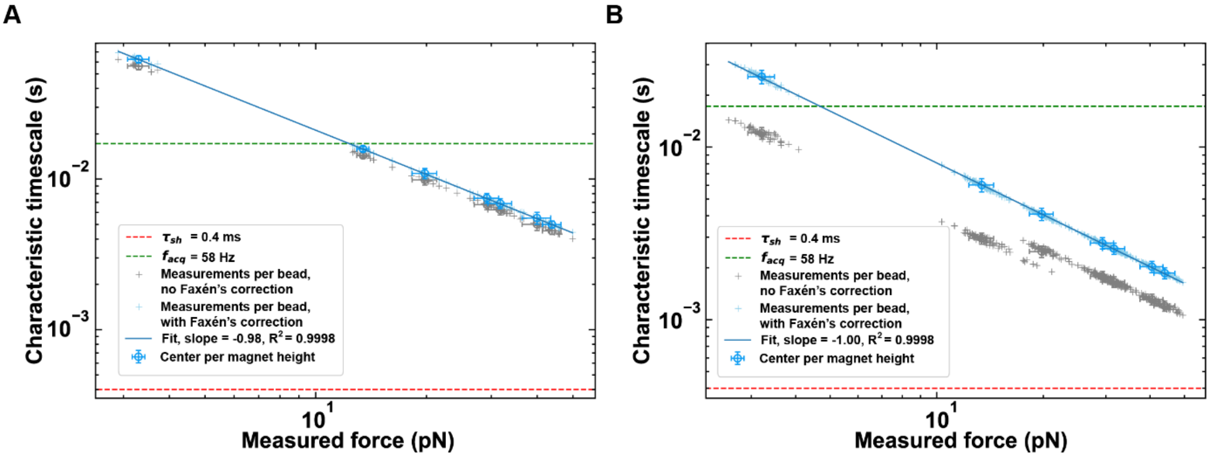


**Figure S4: Characteristic timescales of tethered magnetic beads for the long-pendulum axis fluctuations as a function of force.** The characteristic timescales (s) on the long pendulum axis for multiple beads tethered by **(A)** 20.6 kbp DNA or **(B)** short RNA hairpins (~1.6 kbp handles) versus measured force (pN) using the long pendulum theory. The values for single measurements per bead are indicated as light blue and grey crosses with or without correction for the parallel motion close to the surface in the drag coefficient. The mean and one standard deviation of the values per magnet height with or without Faxén's correction are indicated as blue or grey circles with error bars, respectively. The blue solid line represents a fit to the data using **Eq. 7**. The characteristic timescales need to be higher than the shutter time $\tau_{\rm sh}$ = 0.4 ms (red dashed line) for good force measurements. The acquisition rate $f_{\rm acq}$ = 58 Hz is also indicated.

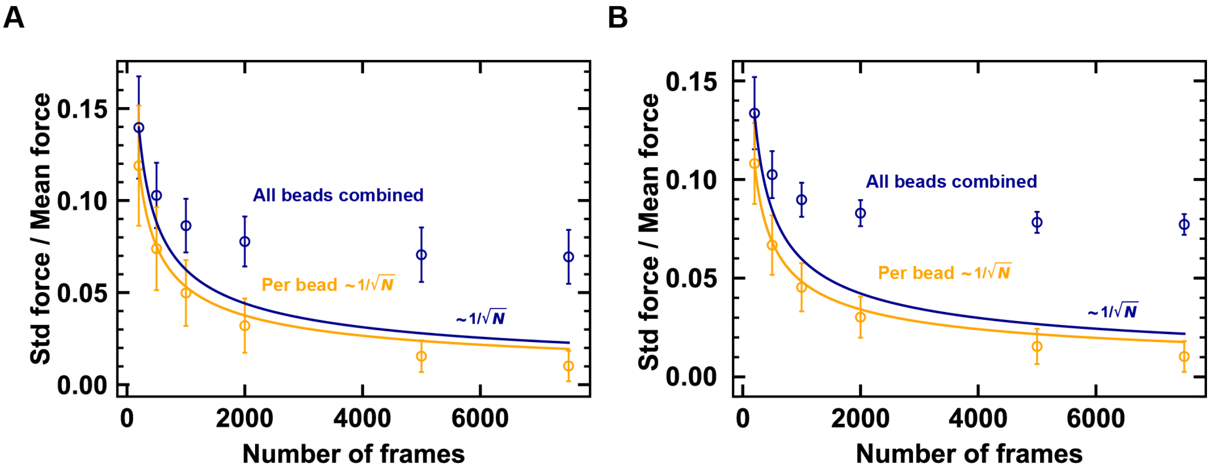


**Figure S5: The relative standard deviation of the forces measured per bead is limited by the statistical error and not the bead-to-bead difference.** The standard deviation (std) in the force measurement divided by the mean force (relative standard deviation) versus the number of uncorrelated frames measured for all bead combined (blue) compared to the forces measured for single beads (orange) for **(A)** long DNA tethers (20.6 kb) and **(B)** RNA hairpins (~1.6 kb handles) compared. The mean (circles) and standard deviation in both direction (error bars) for the RSD on the force measurements are shown compared to the decrease from the statistical error $1/\sqrt{N}$ (lines).

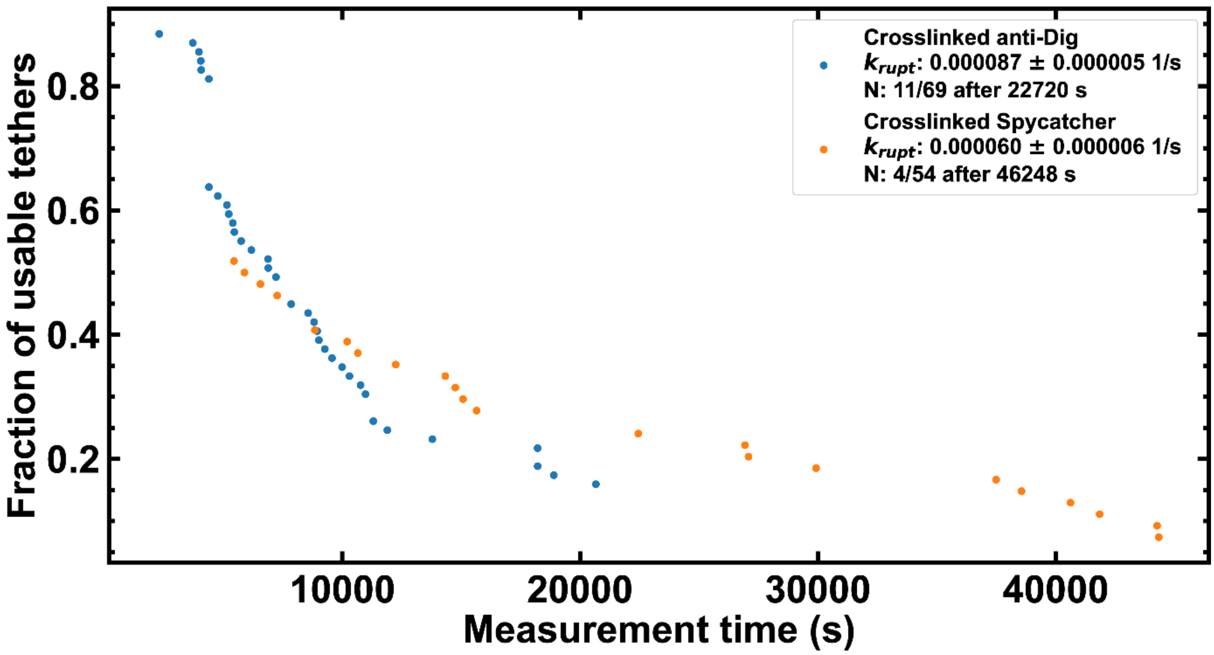


**Figure S6: Comparison of the fraction of usable tethers versus measurement time for different attachment strategies**. The fraction of usable tethers was determined as the number of tethers that showed the expected tether response (**Figure 2B**) divided by the total number of tethers that showed the expected tether response at least in one of the first cycles. Attachment of the DNA hairpin using the Dig-anti-Dig binding on a nitrocellulose surface crosslinked with glutaraldehyde (blue dots) and compared to the attachment using SpyTag-SpyCatcher on a nitrocellulose surface crosslinked with glutaraldehyde (orange dots). The rupture rates of the surface attachment $k_{\mathrm{rupt}}$ are calculated from 100 bootstrap fits.

**Table S1: DNA and RNA hairpins used during the study. (A)** DNA hairpin with MTERF1 binding site, biotin and digoxigenin handles. **(B)** DNA hairpin with MTERF1 binding site, biotin and SpyTag handles. **(C)** RNA hairpin with biotin and digoxigenin handles.

| **Construct** | **Sequence** |
|---|---|
| **DNA hairpin with Digoxigenin handle** | |
| **A** 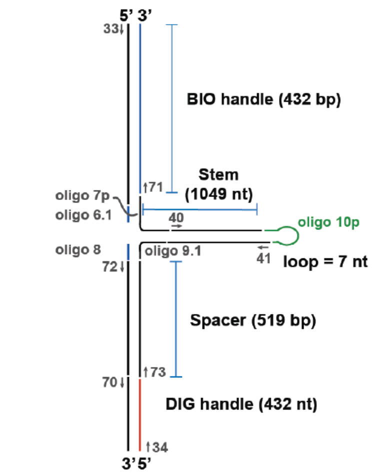  | Oligo 6.1: GCAAAAGTCATTCTGAGAATAGTGTATGCGGCGACCGAGTTG |
| | Oligo 7p: ACCGCAGTACAATCTGCTCTGATGCCGCATGTCTGCACGCTATGTAGAACCAACT |
| | CGGTCGCCGCATACACTATTCTCAGAATGACTT |
| | Oligo 8: TCAATAGATGTGCTGCCCCTCAGTCCGTTGATACCTACTTGCT |
| | Oligo 9.1: TCAAAGCAAGTAGGTATCAACGGACTGAGGGGCAGCACATCTATTGATTTTTTTTTTTTTTTGT |
| | TCTACATAGCGTGCAGACATGCGGCATCAGAGCAGATTGTACTG |
| | Oligo 10p: ATACAGCTCGCCGAGGCGAGAAATCGCCTCGGCGAGCT |
| | Primer 33: AAAAAAGCTTGGAACCAAAGGATATTCAGACGCG |
| | Primer 34: AAAAGGATCCCGTGATGACCTCATTA |
| | Primer 40: CGACATACGGTTGAGACCG |
| | Primer 41: GCGCGGTCTCGGTATTATC |
| | Primer 70: AAAACCTAAGAGACCGGAACCAAAGGATATTCAGACG |
| | Primer 71: AAAAGGTCTCATTGCGGATCCCGTGATGACCTC |
| | Primer 72: AAAAGGTCTCATTGAAACCGGAGTGATGTCGCG |
| | Primer 73: AAAAGGTCTCTTAGGCTTCGCCAGACGGCAT |
| | pMTM plasmid for stem (with or without mutation in the binding site) |
| | WT binding site: 5'-TGTTAAGATGGCAGAGCCCGGTAATCGC-3' |
| | Mutation A(T)3243G(C) in binding site: 5'-TGTTAAGATGGCAG**G**GCCCGGTAATCGC-3' |
| | Mutation G(C)3249A(T) in binding site: 5'-TGTTAAGATGGCAGAGCCCG**A**TAATCGC-3' |
| | Lambda DNA for handles and spacers |
| **DNA hairpin with SpyTag handle** | |
| **B** 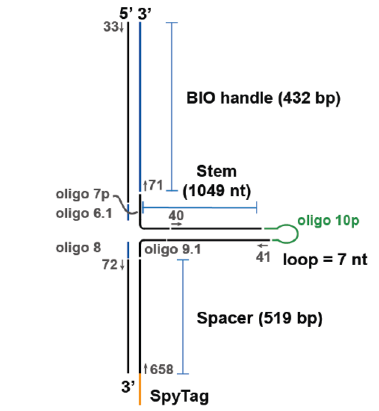  | Oligo's from (A) except for primers 34, 70, and 73 |
| | (*SpyTag-*)Primer 658: *RGVPHIVMVDAYKRYKC* -CTTCGCCAGACGGCATTTAAAGGTG |
| | pMTM plasmid for stem |
| | Lambda DNA for handles and spacers |
| **RNA hairpin with digoxigenin handle** | |
| **C** 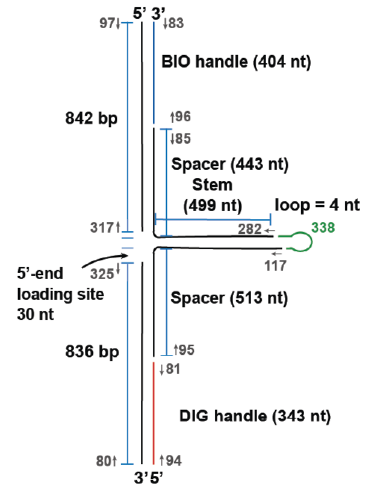  | Primer # 80 - CCTCACTTCTGCTATTTCGC |
| | Primer # 81 - GCGACTTAGCTGAGGCC |
| | Primer # 83 - GGAACCAAAGGATATTCAGACG |
| | Primer # 85 - GGTGCCCACAGAACGTC |
| | Primer # 94 - TAATACGACTCACTATAGGCCTCACTTCTGCTATTTCGC |
| | Primer # 95 - TAATACGACTCACTATAGGTCTTCGCCAGACGGCATTTA |
| | Primer # 96 - TAATACGACTCACTATAGGGCAGGCAAGTCCGATTTTTTG |
| | Primer # 97 - TAATACGACTCACTATAGGGGAACCAAAGGATATTCAGACG |
| | Primer # 117 - TGGATCCGTGGGCGC |
| | Primer # 282 - TAATACGACTCACTATAGTCATGCTCCCATCTTATGG |
| | Primer # 317 - GCCAACTCGGTCGCGTC |
| | Primer # 325 - TAATACGACTCACTATAGGCTGACGTTCTATGCAG |
| | RNA Oligo # 338 - GUAGUGAUUAAAAAUAAUCACUACUGGAUCCGUGGGCGCAGCGGAGAAGAA |
| | pMK-RQ (ColE1 ori, KanR) plasmid (ThermoFisher) with a palindromic sequence |

**Table S2: Fit parameters of single exponential fits to unlocking time distributions for each experimental condition.** The number of traces that did not show an unlocking event within the measurement time $T_{\mathrm{measure}}$ ($N_{\mathrm{cut}}$) relative to the number of measured unlocking events $N_{\mathrm{unlock}}$ are used to correct the unlocking rate $k_{\mathrm{unlock}}$ on long timescales and $t_{\mathrm{cut}}$ is the cut-off on short timescales. (*) Unlocking times from beads experiencing the same calibrated force were combined to build the distribution.

| Measurement | Binding site orientation | Mutation binding site | Mutation MTERF1 | Avg force (pN) | $N_{\mathrm{unlock}}$ | $N_{\mathrm{cut}}$ | $T_{\mathrm{measure}}\ (s)$ | $t_{\mathrm{cut}}\ (s)$ | $k_{\mathrm{unlock}}\ (1/s)$ |
|---|---|---|---|---|---|---|---|---|---|
| WT site; WT MTERF1; perm. | permissive | - | - | 18 | 3813 | 0 | 60 | 0.3 | 0.43 ± 0.01 |
| WT site; WT MTERF1; non-perm. | non-permissive | - | - | 18 | 363 | 0 | 1200 | 0.3 | 0.010 ± 0.001 |
| WT site; WT MTERF1; non-perm. | non-permissive | - | - | 19 | 671 | 1 | 900 | 0.3 | 0.013 ± 0.001 |
| AT3243GC site; WT MTERF1; non-perm. | non-permissive | AT3243GC | - | 19 | 1133 | 1 | 300 | 0.3 | 0.047 ± 0.002 |
| WT site; R162A MTERF1; non-perm. | non-permissive | - | R162A | 19 | 667 | 10 | 180 | 0.3 | 0.038 ± 0.002 |
| AT3243GC site; R162A MTERF1; non-perm. | non-permissive | AT3243GC | R162A | 19 | 118 | 0 | 120 | 0.3 | 0.15 ± 0.03 |
| GC3249AT site; WT MTERF1; non-perm. | non-permissive | GC3249AT | - | 19 | 229 | 0 | 480 | 0.3 | 0.023 ± 0.002 |
| WT site; WT MTERF1; non-perm. | non-permissive | - | - | 21 | 664 | 0 | 240 | 0.1 | 0.060 ± 0.005 |
| WT site; WT MTERF1; non-perm. | non-permissive | - | - | 22 | 657 | 0 | 240 | 0.1 | 0.15 ± 0.01 |
| WT site; WT MTERF1; non-perm.* | non-permissive | - | - | 21.2 ± 0.2 | 128 | 0 | 240 | 0.1 | 0.074 ± 0.008 |
